# Reproduction has different costs for immunity and parasitism in a wild mammal

**DOI:** 10.1101/472597

**Authors:** Gregory F. Albery, Kathryn A. Watt, Rosie Keith, Sean Morris, Alison Morris, Fiona Kenyon, Daniel H. Nussey, Josephine M. Pemberton

## Abstract

1. Life history theory predicts that reproductive investment draws resources away from immunity, resulting in increased parasitism. However, studies of reproductive tradeoffs rarely examine multiple measures of reproduction, immunity, and parasitism. It is therefore unclear whether the immune costs of reproductive traits correlate with their resource costs, and whether increased parasitism emerges from weaker immunity.

2. We examined these relationships in wild female red deer (*Cervus elaphus*) with variable reproductive investment and longitudinal data on mucosal antibody levels and helminth parasitism. We noninvasively collected faecal samples, counting propagules of strongyle nematodes (order: Strongylida), the common liver fluke *Fasciola hepatica* and the red deer tissue nematode *Elaphostrongylus cervi*. We also quantified both total and anti-strongyle mucosal IgA to measure general and specific immune investment.

3. Contrary to our predictions, we found that gestation was associated with decreased total IgA but with no increase in parasitism. Meanwhile, the considerable resource demand of lactation had no further immune cost but was associated with higher counts of strongyle nematodes and *Elaphostrongylus cervi*. These contrasting costs arose despite a negative correlation between antibodies and strongyle count, which implied that IgA was indicative of protective immunity.

4. Our findings suggest that processes other than classical resource allocation tradeoffs are involved in mediating observed relationships between reproduction, immunity, and parasitism in wild mammals. In particular, reproduction-immunity tradeoffs may result from hormonal regulation or maternal antibody transfer, with parasitism increasing as a result of increased exposure arising from resource acquisition constraints. We advocate careful consideration of resource-independent mechanistic links and measurement of both immunity and parasitism when investigating reproductive costs.

## Introduction

Costly traits are central to the fields of life history theory and ecoimmunology. Tradeoffs arising between reproductive investment and other aspects of life history are a fundamental prediction of the former (Harshman & Zera, 2007; Stearns, 1989; Williams, 1966), while the latter examines the ecology of costly immune responses (Graham et al., 2011; Sheldon & Verhulst, 1996). Because reproduction and immunity compete for host resources, in resource-limited environments, animals that invest in reproduction should have fewer resources to allocate to immune defences (Deerenberg, Arpanius, Daan, & Bos, 1997; French, Denardo, & Moore, 2007; Sheldon & Verhulst, 1996). If immunity is protective, these individuals will experience higher parasitism as a result. Intuitively, traits with higher resource demands should result in the diversion of more resources away from immunity, leading to higher parasite burdens. However, recent advances have demonstrated that the interrelationships between host resources, immunity, and parasitism can be unexpectedly complex (Cressler, Nelson, Day, & Mccauley, 2014). Few studies in wild mammals have examined tradeoffs with multiple reproductive traits, so it is unclear whether different components of reproduction have different costs for immune defence, and whether their costs are proportional to their resource demand. Furthermore, studies of reproductive tradeoffs rarely quantify both immunity and parasitism to examine the importance of susceptibility versus exposure in driving higher parasite intensities in reproductive females (Bradley & Jackson, 2008; Knowles, Nakagawa, & Sheldon, 2009). Here, we examine the partitioning of reproductive costs for multiple measures of immunity and parasitism to investigate the possible mechanisms governing a reproduction-immunity-parasitism tradeoff in a wild mammal.

Mammalian reproduction is a complex, multi-stage process, featuring extensive maternal investment which varies in intensity through the reproductive period (Langer, 2008; Maestripieri & Mateo, 2009). As such, different components of reproduction vary substantially in their resource and fitness costs. In particular, lactation is a highly energetically demanding process which carries costs for immunity, parasitism or fitness in a range of mammals (Beasley, Kahn, & Windon, 2010; Christe, Arlettaz, & Vogel, 2000; Clutton-Brock, Albon, & Guinness, 1989; Froy, Walling, Pemberton, Clutton-brock, & Kruuk, 2016; Jones, Sakkas, Houdijk, Knox, & Kyriazakis, 2012; Rödel, Zapka, Stefanski, & von Holst, 2016; Woodroffe & Macdonald, 1995). Meanwhile, only one of these studies uncovered an immunological cost of gestation (Christe et al., 2000), which generally requires fewer resources than does lactation (Clutton-Brock et al., 1989). However, although experimentally modifying resource availability can affect the severity of reproduction-immunity tradeoffs (French et al., 2007; Jones et al., 2012), this is not always the case (Stahlschmidt, Rollinson, Acker, & Adamo, 2013). Similarly, studies in birds have questioned whether the energetic costs of immunity are sufficient to drive tradeoffs (Eraud, Duriez, Chastel, & Faivre, 2005; Svensson, Råberg, Koch, & Hasselquist, 1998). Such findings imply that reproduction-immunity tradeoffs can be linked mechanistically as well as through resource reallocation. Potential such links include production of reactive oxygen species, reduction in immunologically active fat stores, or resource-independent hormonal regulation (Speakman, 2008; Svensson et al., 1998).

Different components of mammalian reproduction can have qualitatively different effects on host immunity as well as varying quantitatively in terms of their resource demand. For example, pregnancy necessitates modulation of the immune system to avoid mounting an immune response to the developing foetus, which will directly affect anti-parasite defence (Weinberg, 1984, 1987). Similarly, lactation draws immune molecules away from the mother for transfer to offspring, reducing their availability for the mother’s own defence (Grindstaff, Brodie, & Ketterson, 2003; Hasselquist & Nilsson, 2009). Reproduction also induces a suite of physiological and behavioural changes which will affect susceptibility and exposure to parasites indirectly: for example, it has been suggested that bats compensate for the energetic demand of lactation by reducing costly grooming behaviour, with ectoparasite burden increasing as a result (Speakman, 2008). It is unclear how such mechanistic links between components of reproduction and immunity interact with resource allocation to influence immune defence and parasite intensity in wild mammals.

The wild red deer (*Cervus elaphus*) in the North block of the Isle of Rum exhibit a well-studied life history tradeoff, in which reproduction substantially decreases the mother’s probability of overwinter survival and reproduction the following year (Clutton-Brock et al., 1989; Froy et al., 2016). However, not all components of reproduction are equally costly: gestation has a negligible detectable fitness cost compared to that of lactation (Clutton-Brock et al., 1989). Moreover, while giving birth late and caring for a male calf compared to a female calf are associated with decreased maternal fitness, their effects are small compared to the cost of lactation itself (Froy et al., 2016). The study population has a high prevalence of gastrointestinal helminths, and parasite burdens can be quantified noninvasively through faecal propagule counts (Albery et al., 2018). Mucosal antibodies, and especially the IgA isotype, are important effectors of ruminant adaptive immunity to gut helminths (Butler, 1969; McRae, Stear, Good, & Keane, 2015). Mucosal IgA can be quantified in wild ruminant faeces, correlating positively with the same isotype measured in plasma or serum and negatively with helminth faecal egg counts (Watt, Nussey, Maclellan, Pilkington, & McNeilly, 2016).

In this study, we measured both total and helminth-specific mucosal IgA and propagule counts of multiple helminth species, using faecal samples collected from the Isle of Rum study population. We quantified the associations between several reproductive traits of known fitness cost and subsequent measures of immunity and parasitism. We also examined covariance between IgA and parasites to discern whether increased IgA was associated with decreased parasite intensity independently of shared reproductive and seasonal effects, implicating IgA as an indicator of protective immunity. We predicted that reproductive investment would be associated with reduced antibody levels and increased parasite counts, and that aspects of reproduction previously found to be more costly for fitness, especially lactation, should likewise be more costly in terms of both immunity and parasitism. Furthermore, providing parasitism is mediated by immune susceptibility, aspects of reproduction that are costly for immunity should have similar costs in terms of parasitism.

## Methods

### Study system, sampling and parasitology

The study population is located in the North block of the Isle of Rum National Nature Reserve in the Inner Hebrides, Scotland (57°N 6°20’W). The resident population comprises ∼350 animals at any one time, and is regularly censused to keep track of each individual and its life history. See Clutton-Brock *et al*. (1982) for a full summary of the project and the deer reproductive cycle. Briefly, the deer mate in September and October and give birth in May-June, after an approximately 235 day gestation. Females do not reproduce every year, and produce a maximum of one calf per year. During the calving season, daily monitoring of pregnant females enables the recording of precise birth dates. Neonates are caught, sexed, weighed and individually marked, enabling life-long individual identification. Calves are dependent on their mothers for much of their first year. Regular population censusing throughout the year and winter mortality searches allow dates of death to be reliably assigned to the nearest month for the vast majority of individuals. Most calf deaths occur either within the first few weeks of life, or in the following winter ∼6-9 months later. Females that successfully raise a calf to the age of one, or that lose the calf in its first winter, have lower rates of overwinter survival and reproduction the following year compared to those that do not reproduce that year or that lose their calf in the summer (Clutton-Brock et al., 1989; Froy et al., 2016). Many calves die over the winter, but the mothers of these calves have paid the cost of lactation associated with feeding them until the winter, whether or not the calf survive. Therefore these females are treated as a single category here (Clutton-Brock et al., 1989; Froy et al., 2016).

We collected faecal samples from female deer across the annual reproductive cycle. As a “deer year” runs from May to April, this study examines the effects of reproduction over a year, beginning in May, on egg counts and antibody levels until the following April. A description of the sample collection procedure can be found in Albery *et al*. (2018). Sampling occurred over seven two-week sampling trips spanning April 2016-April 2018, in August (“summer”), November (“autumn”) and April (“spring”). Note that our dataset included an April sampling trip from the deer reproductive cycle starting May 2015, without an accompanying August and November trip from this reproductive cycle. Figure 1 illustrates how sampling relates to different aspects of reproductive investment by female deer across the annual cycle. We classify a female’s reproductive status for a given year as “No Calf”, “Calf Died” and “Calf Survived” (see Figure 1). “No Calf” samples were collected from females that did not reproduce in the calving season preceding the sampling trip; “Calf Died” samples were collected from females that gave birth to a calf in the preceding calving season which died before October 1^st^ of that year; and “Calf Survived” samples were collected from females that gave birth to a calf in the preceding calving season which survived past October 1^st^ of that year. We excluded females that were reproducing for the first time from our analyses, as their reproductive success is heavily confounded with their young age (mean age 4.21 years). In addition, females may or may not become pregnant during the autumn rut. Samples were therefore assigned a pregnancy status, beginning in November, based on whether or not the female gave birth to a calf in the following spring (Figure 1).

**Figure 1:**
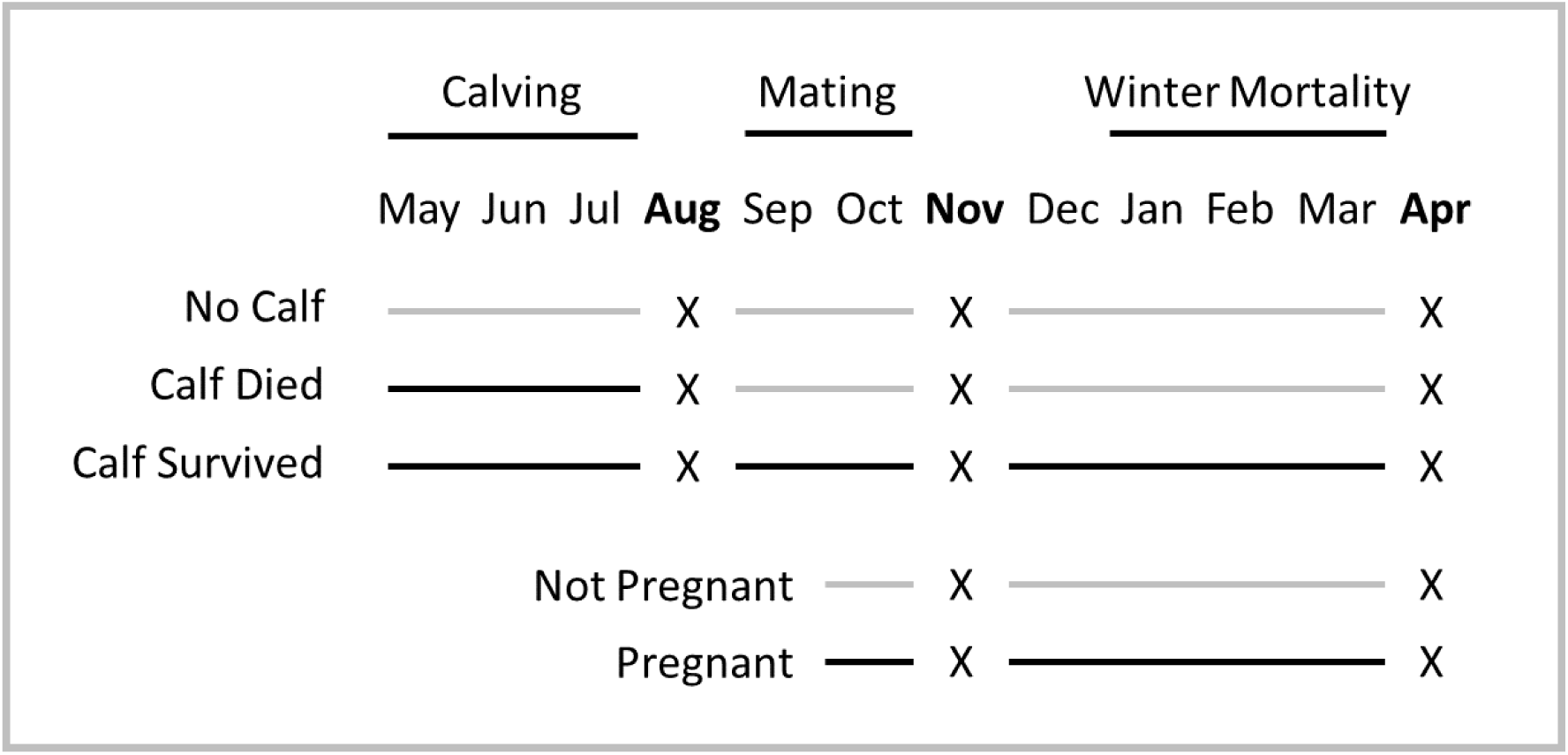
The faecal sampling regime in the context of a deer reproductive cycle (“deer year”). A cross (X) represents a two-week sampling trip. The deer year begins on May 1^st^when calving begins; individuals are assigned to one of three reproductive status categories (top three lines) according to the birth and survival of their calf over the course of the following year. Individuals are also assigned a pregnancy status in November and April based on reproduction in the following calving season (bottom two lines). Black lines represent periods in which the calf is living or the female is pregnant; grey lines represent periods in which the calf is dead or non-existent or the female is not pregnant.

In total 837 faecal samples were collected noninvasively from 140 mature females. In the evening after collection, samples were homogenised by hand and subsampled, with 1-15g frozen at −20°C for antibody quantification and the remainder refrigerated at 4°C for parasitological analysis. Subsamples were transferred to Edinburgh at these temperatures. Parasite propagule counts were carried out as previously described, without correcting for dry weight, and included counts of strongyle nematodes, the common liver fluke *Fasciola hepatica* and the tissue nematode *Elaphostrongylus cervi* (Albery et al., 2018). Final sample sizes for each variable are displayed in Table SI1.

### Antibody extraction and quantification

Faecal antibodies were quantified using a protocol modified from Watt *et al*. (2016). Faecal matter was stored at −20°C until extraction. Extractions occurred either in January-March 2017 (session “A”, samples collected April-November 2016; N=132), January 2018 (“B”, samples collected April-November 2016; N=212) or within the sampling trip (“C”, samples collected April 2017-April 2018, N=460). 0.6g (+/- 0.005g) of the homogenate was weighed out into an Eppendorf tube and mixed thoroughly with 0.9ml of protease inhibitor solution (cOmplete ™ Mini Protease Inhibitor Cocktail tablets, Roche, Basel, Switzerland; 1 tablet mixed with 7ml Phosphate Buffered Saline). The mixture was left to stand for a minimum of 5 minutes to allow the protease to act and then centrifuged at 10,000g for 5 minutes. The supernatant was removed using a micropipette and stored in a separate Eppendorf tube at −20°C until ELISA.

We measured two antibodies by ELISA: total IgA and anti-*Teladorsagia circumcincta* third larval stage IgA (anti-Tc IgA). *T. circumcincta* is an abundant and important sheep strongyle, and anti-Tc IgA correlates negatively with strongyle (order: Strongylida) faecal egg count in wild Soay sheep (Watt et al., 2016). *T. circumcincta* is also present in the Rum deer (unpublished data). ELISA plates were coated the night before using sheep-derived capture antibodies (Bethyl Laboratories, Montgomery, TX) for total IgA and with third larval stage antigen for anti-Tc IgA (Moredun Research Institute, Penicuik, Scotland). For total IgA the samples were diluted in the ratio 1:64; due to lower antibody concentrations undiluted supernatant was used for the anti-Tc IgA assay. After this stage, the ELISA protocol was carried out as described in Watt *et al*. (2016). The total IgA dilution was selected by carrying out serial dilutions on a set of samples and selecting the dilution at which different concentrations of antibodies were deemed to have the widest spread of optical densities. ELISA readings diluted linearly as expected. Samples were corrected using controls according to the calculation: Final OD=(sample OD-mean plate negative OD)/(mean plate positive OD- mean plate negative OD). All samples were run on duplicate plates, which were highly correlated (R=0.98 across all duplicates). The mean value for the two duplicates was taken for each sample and used for analysis.

### Statistical analysis

We used four sets of Generalised Linear Mixed Models (GLMMs) to test how reproductive traits were associated with antibody levels and parasite intensity. Analyses were carried out in R version 3.5.0 (R Core Team 2018) with the package MCMCglmm (Hadfield, 2010). All models were run for 2.6×10^6^ iterations with a 2000 iteration thinning interval and a 6×10^5^ iteration burn in period. P_MCMC_ values for differences between factor categories were calculated using the proportional overlap of estimates’ posterior distributions, divided by half the number of iterations.

### Full models

We first constructed five univariate GLMMs using the full dataset (837 samples from 140 individuals). Three models used an overdispersed Poisson distribution, with strongyle, *F. hepatica* and *E. cervi* intensity as response variables. Models initially included the following fixed effects, without interactions: Year (factor with three levels representative of the deer reproductive cycle beginning in 2015, 2016 and 2017); Season (factor with three levels: Summer, Autumn and Spring); Age in years (continuous); and Reproductive Status (factor with three levels: No Calf, Calf Died and Calf Survived). Individual identity was fitted as a random effect. All continuous variables except parasite counts were scaled to have a mean of 0 and a standard deviation of 1 before analysis.

The two remaining models examined antibodies as response variables. As mucosal antibodies are vulnerable to degradation by temperature-dependent faecal proteases, we had to account for the extraction session and time to freezing and extraction (Figure SI5-6). There was an uneven distribution of year, season, and status categories across different extraction sessions, so that these variables could not all be fitted in the same model. Therefore, to control for collection factors and quantify reproductive status effects conservatively we first transformed antibody levels to approximate normality (square-root transform for total IgA and cube-root transform for anti-Tc IgA), and fitted a linear model with fixed effects including extraction sessions performed at different times (factor with three levels); day of collection within a sampling trip (continuous integers, range 0-11); time elapsed from sample collection until freezing (continuous, in hours). The scaled residual values from these models (mean=0, SD=1) were used as the response variables in two Gaussian GLMMs with the same fixed and random effects as the parasite GLMMs.

Previous work on the Rum deer revealed extensive seasonal fluctuations in parasite count (Albery et al., 2018). We therefore followed up the above five models by fitting a season by reproductive status interaction in order to investigate whether the effects of reproductive status varied by season. Each model was compared with and without this interaction to investigate whether it affected Deviation Information Criterion (DIC) values as a measure of model fit (threshold values for distinguishing between models ΔDIC=2) or changed model estimates.

### Pregnancy models

Pregnancy may directly affect immunity, and effects attributed to reproductive status could be due to correlated variation in pregnancy status over the sampling year. To check this we ran a second set of models investigating the role of pregnancy status. This used a subset of samples collected in November and April (518 samples from 122 individuals), as mating occurs in the early autumn and females could not be pregnant in August. These five models featured the same explanatory variables as the full status models, with only two levels in the season category (Autumn and Spring), and with Pregnancy included as a binary variable. We compared these models with and without the pregnancy term as a fixed effect to investigate whether its inclusion changed reproductive status effect sizes or affected model fit.

### Calving trait models

We next used a restricted dataset consisting of individuals that had given birth in the year of sampling (571 samples from 116 individuals) to investigate whether specific traits associated with a calving event influenced antibody levels and parasitism. We fitted the same fixed and random effects as the full model set, but with only two factor levels in the reproductive status category (Calf Died and Calf Survived), and including several variables relating to each birth: Parturition Date (continuous, centred around median birth date that year); Birth Weight (continuous, in kilograms, calculated from a projection using capture weight and age in hours, slope 0.01696 kg/h); Calf Sex (Female or Male).

### Multivariate model

Multivariate mixed-effects models can be used to investigate covariance between measures of immunity and parasitism, while accounting for fixed effects. To test whether antibodies and parasites were correlated we fitted a model with strongyles, *E. cervi*, total IgA and anti-Tc IgA as response variables, with the same fixed effects as the full univariate models. Unstructured variance/covariance matrices were fitted for random and error terms, allowing estimation of the phenotypic correlations between the response variables. Phenotypic covariance between two response variables A and B (Cov_phenotypicA,B_) is calculated using the random (G) and residual (R) variance structure of the model, with the formula Cov_phenotypicA,B_=Cov_IndividualA,B_+Cov_residualA,B_. Phenotypic correlation between two response variables (r_phenotypicA,B_) was calculated by dividing the phenotypic covariance by the square root of the sum of the variance in both response variables: r_phenotypicA,B_=Cov_phenotypicA,B_/(V_phenotypeA_+V_phenotypeB_)^0.5^. P_MCMC_ values for correlations were calculated using the posterior distributions, dividing the number of iterations overlapping with zero by half the total number of iterations.

## Results

Reproductive investment was strongly associated with both lower antibody levels and increased parasite counts, but patterns differed considerably between different response variables (Figure 2, SI1). Compared to “No Calf” individuals, “Calf Survived” status was associated with higher intensity strongyle (P_MCMC_<0.001) and *E. cervi* infection (P_MCMC_=0.01), and with lower total IgA (P_MCMC_=0.016) and anti-Tc IgA levels (P_MCMC_<0.001). “Calf Survived” females also had higher parasite counts than “Calf Died” individuals (P_MCMC_<0.001 for strongyles and *E. cervi*), but these reproductive status categories did not differ in total IgA (P_MCMC_=0.502) or anti-Tc IgA (P_MCMC_=0.336; Figure 2-3). “Calf Died” individuals did not differ from “No Calf” females in strongyle, *E. cervi* or anti-Tc IgA levels (Figure 2) but had lower total IgA levels (P_MCMC_=0.018). That is, “Calf Died” individuals had lower total IgA than “No Calf” females, but with similar parasite intensities, while “Calf Survived” individuals had the same antibody levels as “Calf Died” individuals, but with increased parasite intensities. *F. hepatica* was not associated with reproductive status, but decreased with age (P_MCMC_=0.004) as did *E. cervi* (P_MCMC_<0.001; Figure SI1, SI7).

**Figure 2:**
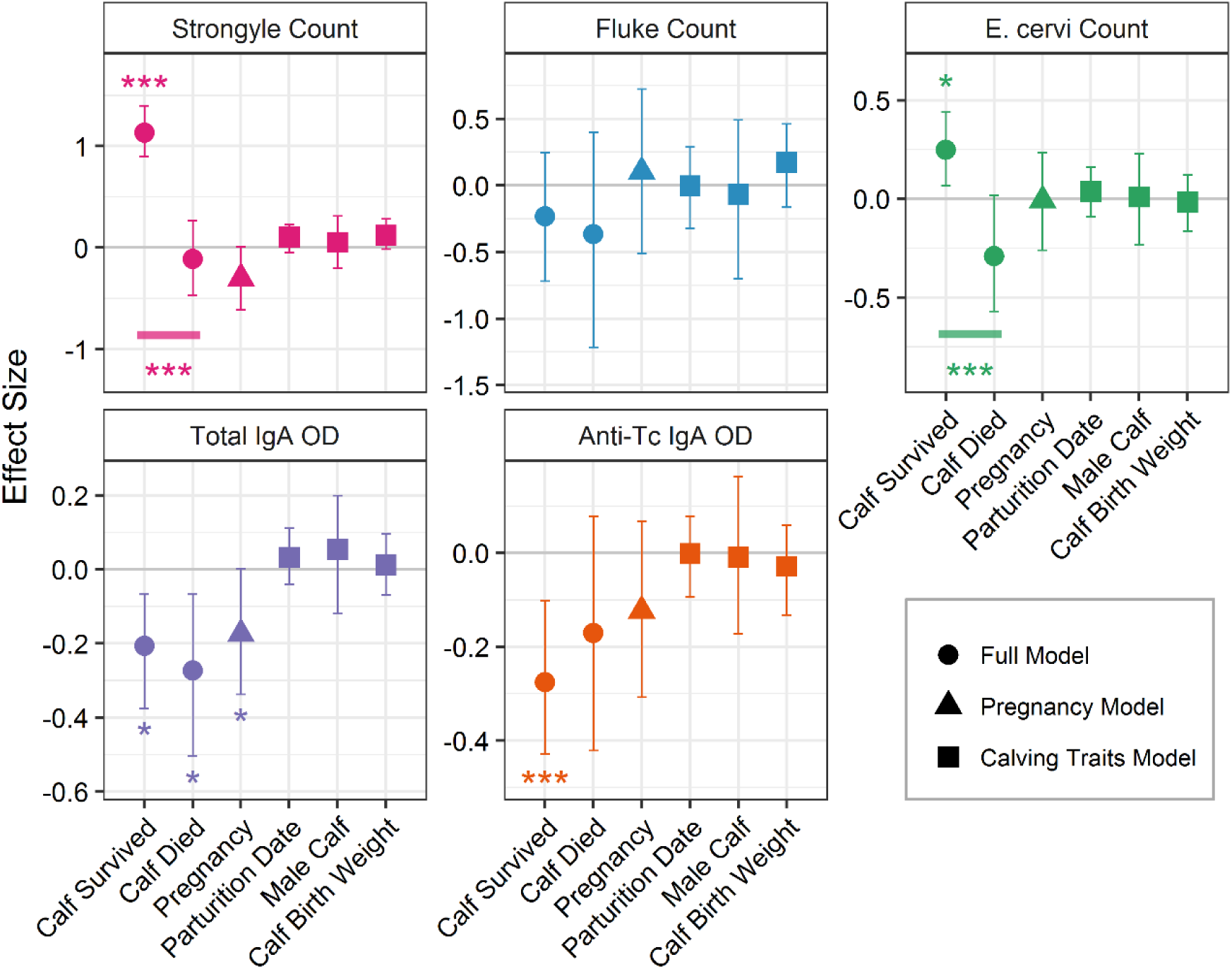
Model outputs depicting the effects of reproductive traits, derived from all three univariate model sets. Points and error bars represent model estimates and 95% credibility estimates. Effect sizes for categorical variables (status, pregnancy and calf sex) denote differences from the first (absent) category of each, contained in the intercept (“No Calf”, “Not Pregnant” and “Female Calf” respectively). Effect sizes for continuous variables (parturition date and calf birth weight) represent the change in the response variable associated with a change of one standard deviation of the explanatory variable. Asterisks represent significant differences derived from MCMCglmm posterior distribution overlaps: ***, ** and * denote P<0.001, P<0.01 and P<0.05 respectively. Bars denote differences between status categories.

Strongyles and both antibodies all exhibited the same seasonality, peaking in the spring and being lowest in the autumn, with the summer intermediate (Figure 3, all differences P_MCMC_<0.001). *F. hepatica* was higher in the spring than in the summer or autumn (P_MCMC_<0.034), and *E. cervi* was lowest in the summer, with the autumn intermediate (P_MCMC_<0.001). There was also some between-year variation: strongyle levels increased between 2015 and 2016, and again in 2017 (all P_MCMC_<0.001), while total IgA levels decreased in 2017 compared to 2015 and 2016 (P_MCMC_<0.024). Anti-Tc IgA was also lower in 2017 than 2016 (P_MCMC_<0.001). Inclusion of season-by-status interactions improved strongyle model fit (ΔDIC=-3.79), but did not improve the fit of any other models (ΔDIC<2). Fixed status effects remained largely unchanged in magnitude or significance, suggesting that the observed associations with reproductive status were consistent across seasons (Figure 3). All interaction terms implied an attenuation of reproductive status effects from summer through winter to spring, rather than any major qualitative change in this association (Figure 3). Both “Calf Died” and “Calf Survived” females had increased antibody levels and decreased parasite intensities relative to “No Calf” females over this period. See Figure SI2 for a comparison of the full model estimates and DIC changes when a season-by-status interaction was included.

**Figure 3:**
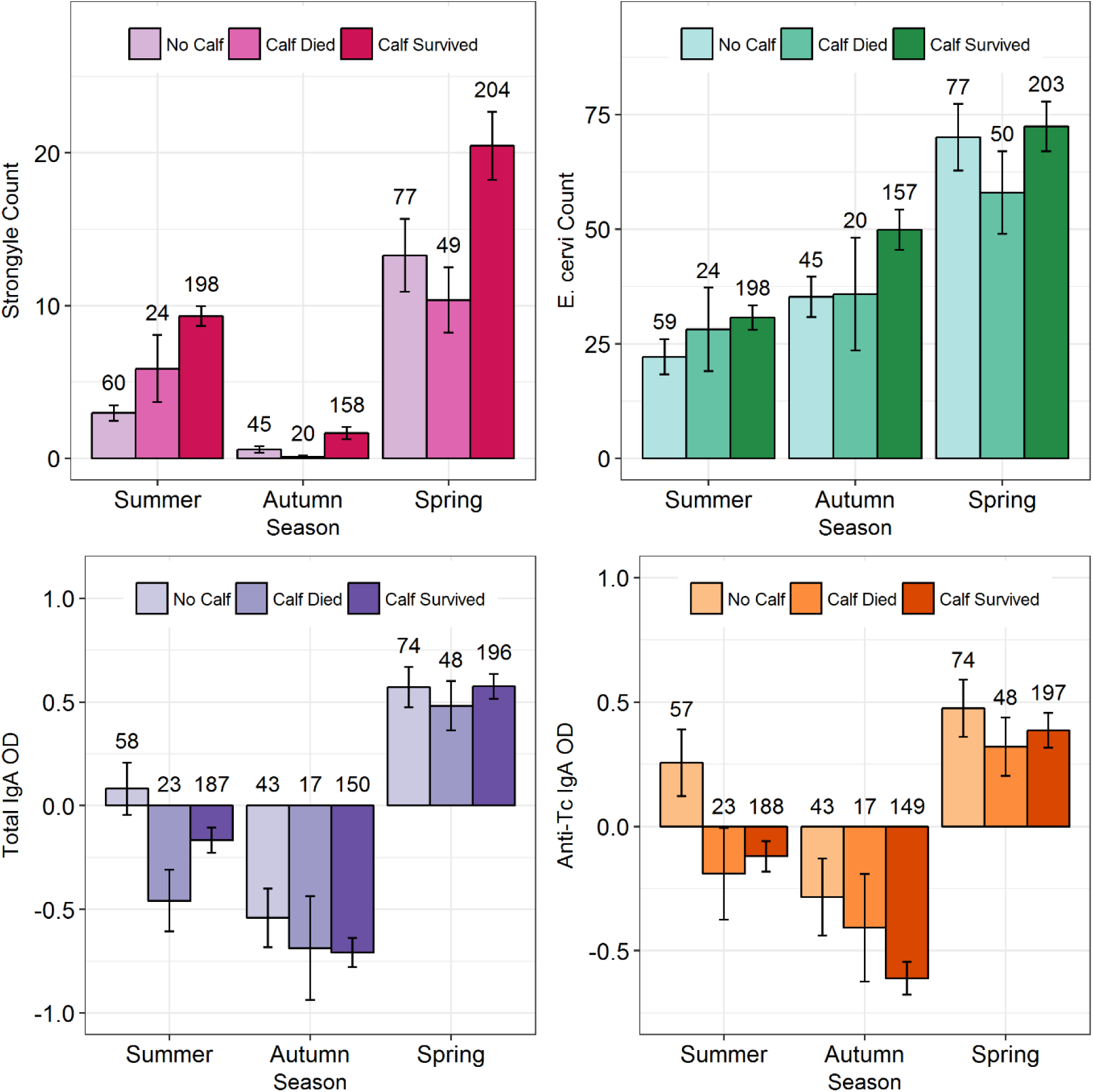
Bar charts displaying raw mean (+/- SE) parasite counts and antibody levels of each reproductive status category in each season. Antibody measures were taken from the residuals of a model with square root-transformed (total IgA) or cube root-transformed (anti-Tc IgA) antibody OD as the response variable and including collection variables as fixed effects. Numbers above bars denote sample sizes.

Pregnancy models examining April and November samples revealed marginally lower total IgA in pregnant females (P_MCMC_=0.034, Figure 2, SI1, SI3). Including pregnancy status in our models did not alter the direction or significance of reproductive status effects; in fact, in the case of total IgA and anti-Tc IgA it increased the significance of the “Calf Survived” category’s effect (Figure SI3). It also slightly improved the fit of the total IgA model (ΔDIC=-3.00). No other models were impacted notably by the inclusion of the pregnancy term, although it slightly reduced the effect size of the “Calf Survived” category in influencing strongyle count (Figure SI3). Although the “Calf Died” category was not significant in the total IgA pregnancy model as it was in the full model, the fact that adding and removing pregnancy as a variable had very little effect on the model estimate (Figure SI3) implies that this did not arise from confounding effects of pregnancy.

None of the calving traits modelled (parturition date, calf birth weight or calf sex) were associated with maternal parasite or antibody levels (Figure 2, SI1).

The fixed effects of the multivariate model were very similar to those of the full models (Figure SI4). The raw correlations between the response variables of the model are displayed in Figures 4 and SI8. Phenotypic correlations (R_p_) derived from the variance structure of the multivariate model are as follows. There were strong positive correlations between strongyles and *E. cervi* (R_p_=0.26, P_MCMC_<0.001) and between total and anti-Tc IgA (R_p_=0.424, P_MCMC_<0.001). Strongyle count was also weakly negatively correlated with total IgA (R_p_=- 0.074, P_MCMC_=0.016, Figure SI8) and more strongly with anti-Tc IgA (R_p_=-0.142, P_MCMC_<0.001, Figure 4).

**Figure 4:**
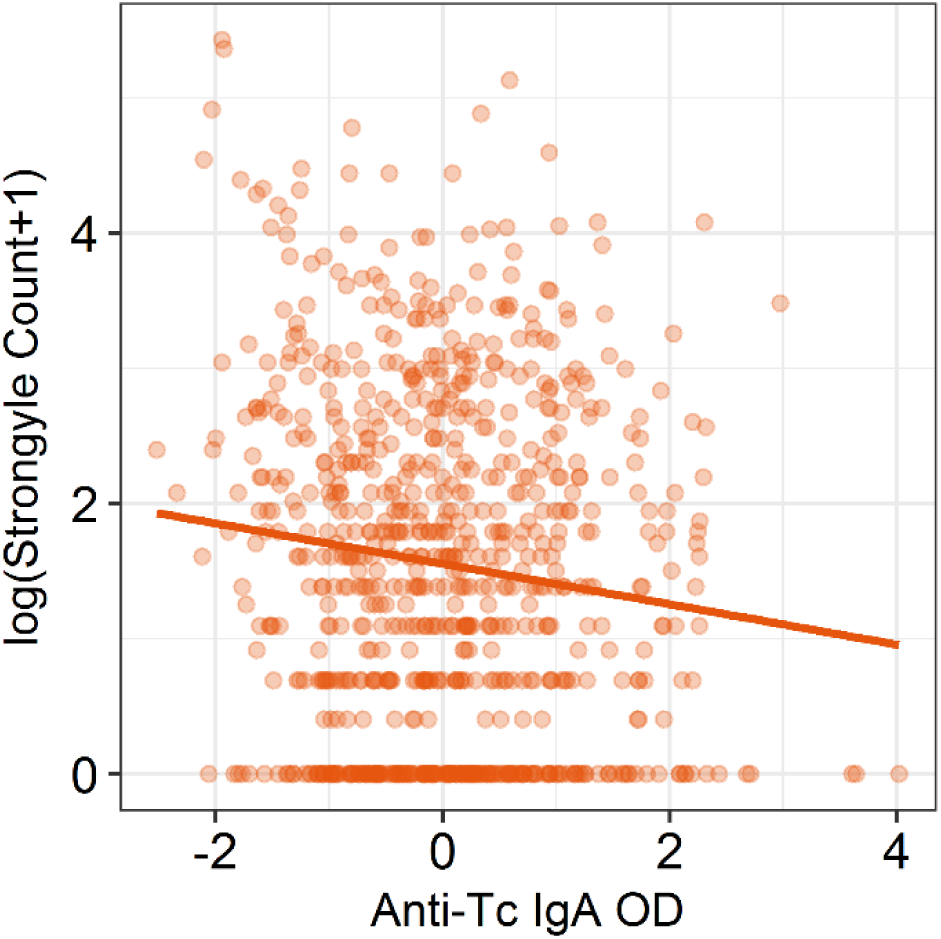
Correlation between anti-*Teladorsagia circumcincta* IgA levels and strongyle count, using the raw data. Individuals with higher anti-Tc IgA had lower strongyle counts (multivariate model phenotypic correlation R_p_=-0.142, P_MCMC_<0.001). The y axis is on the log_e_(count+1) scale to aid interpretation; the x axis data were taken from the residuals of a model with cube root-transformed anti-Tc IgA as the response variable and including collection variables as fixed effects. For this figure, these residuals were centred within sampling trips to have a mean of 0 and a standard deviation of 1 to avoid a positive correlation arising from shared seasonal and annual effects.

## Discussion

Lactation is associated with weaker immunity or increased parasitism in a range of mammals (Festa-Bianchet, 1989; Jones et al., 2012; Rödel et al., 2016; Woodroffe & Macdonald, 1995). In accordance with these studies, we found that lactating females had both decreased antibody levels and increased parasite counts relative to non-reproductive females. In contrast, gestation is rarely found to be costly for immunity or parasitism in mammals (Irvine, Corbishley, Pilkington, & Albon, 2006; Rödel et al., 2016; Woodroffe & Macdonald, 1995), and carries no detectable fitness cost in the Rum red deer (Clutton-Brock et al., 1989). Here, deer that gave birth to a calf that died as a neonate, thereby incurring a limited lactation cost, had lower total IgA levels than non-reproducing females. Gestation therefore carried an immune cost in this study. We predicted that resource depletion incurred through investment in a given reproductive trait would lead to reduced immune investment, and that this would lead to increased parasite intensity (Knowles et al., 2009; Sheldon & Verhulst, 1996). Our results deviated from our expectations in two ways: first, gestation’s long-lasting immune cost was not accompanied by increased parasite intensities. Second, the considerable additional resource investment of prolonged lactation was not associated with additional immune costs relative to gestation, but was instead associated with an increase in parasite intensity. These results have two implications: reproduction-immunity tradeoffs were unlikely to be mediated by simple resource reallocation, and reproduction-parasitism tradeoffs were unlikely to be mediated solely by immunity – despite our observation that higher immune investment was associated with lower parasite counts between individuals (Figure 4, SI8).

If gestation’s lack of detectable fitness cost in our study population (Clutton-Brock et al., 1989) demonstrates a small resource cost, why was gestation associated with reduced total IgA levels, and why did the additional resource cost of lactation not decrease antibody levels further? First, it is possible that reproductive hormones suppress the immune system without being sensitive to resource availability (Foo, Nakagawa, Rhodes, & Simmons, 2017; Svensson et al., 1998). Similarly, gestation may lead to alterations in immune investment and antibody production, so that lower IgA resulted from selective investment in alternative immune cells or functions rather than from lower absolute resource investment in immunity. Reproductive mammals are commonly found to exhibit different (rather than weaker) immunity, but specific patterns of immune prioritisation are unpredictable. For example, reproductive vampire bats (*Desmodus rotundus*) prioritise innate over adaptive immunity (Becker et al., 2018), while reproductive rabbits (*Oryctolagus cuniculus*) exhibit reduced white blood cell counts but stronger humoral immunity (Rödel et al., 2016). Assessing whether reproductive deer invest preferentially in aspects of immunity other than mucosal antibodies would therefore necessitate examining numerous additional immune measures – however, in this study we were restricted to quantifying mucosal antibodies using noninvasive faecal samples as the deer are rarely handled as adults (Clutton-Brock et al., 1982).

Alternatively, gestation and early lactation may necessitate export of IgA from the gut to the blood for transfer to offspring (Jeffcoate et al., 1992; Sheldrake, Husband, Watson, & Cripps, 1984). In ungulates a substantial proportion of maternal antibody transfer occurs via the colostrum in the first few days of life (Hurley & Theil, 2011). It is feasible that this diversion of IgA from the gut occurs around parturition and is detectable for an extended period of time without an underlying resource allocation tradeoff, creating lower IgA levels in all reproductive females regardless of their calf’s survival. The necessity of transferring immune effectors to offspring may therefore be an important obligate mechanism contributing to reduced antibody levels in reproductive wild mammals (Rödel et al., 2016). In a proposed mechanism for the periparturient rise in helminth egg count in domestic sheep, exportation of IgA from the gut around parturition releases helminths from immune control (Jeffcoate et al., 1992). However, in this study, the lower total IgA and intermediate anti-Tc IgA levels in female deer that only paid the cost of gestation were not accompanied by any change in parasitism. This is surprising, given that the results of our multivariate model implied that both IgA measures are representative of increased resistance to strongyles (Figure 4, SI8).

If antibody levels were indicative of investment in protective immunity, how were the deer that paid the immune cost of gestation able to maintain low strongyle and *E. cervi* intensities? Or, what produced the higher parasite counts in lactating individuals? Lactating females’ anti-Tc IgA levels were significantly lower than nonreproductive females’, which could explain their increased parasitism in the absence of a contrast with any other reproductive categories. However, levels of total and anti-Tc IgA in lactating females were not detectably lower than those exhibited by females that paid the cost of gestation (Figure 2). This disparity suggests that additional processes such as exposure were important in driving the high parasite intensities in lactating females (Knowles et al., 2009; Sheldon & Verhulst, 1996). The energetic and resource demand of milk production necessitates substantially increased forage intake and grazing time (Hamel & Côté, 2008, 2009), and may reduce feeding selectivity or the ability to exhibit parasite avoidance behaviours (Hutchings, Judge, Gordon, Athanasiadou, & Kyriazakis, 2006; Speakman, 2008). Thus, lactating females may suffer increased exposure to infective larvae, resulting in higher parasite burdens. This mechanism offers an explanation for our observation that lactation was associated with increased parasite counts, while gestation was not, as individuals that lost their calf as a neonate were not then saddled with a necessity for such high resource acquisition. Based on our results, we suggest that severe effects of mammalian reproduction on parasite infection are partly mediated by exposure as a result of constraints on resource acquisition, foraging selectivity, and antiparasite behaviours, in addition to increased immune susceptibility.

Effects of foraging on exposure can profoundly affect epidemiological dynamics: for example, in the water flea *Daphnia dentifera*, temperature-induced increases in food intake can increase the magnitude of fungal pathogen epidemics (Shocket et al., 2018). Similar processes may act in the deer, if spatiotemporal variation in climatic conditions, deer density, or food abundance modify feeding behaviour or the threat of exposure. In particular, strongyle and *E. cervi* parasitism will be further exacerbated in years and areas of the study system where deer density is high and food availability is low (Wilson, Grenfell, Pilkington, Boyd, & Gulland, 2004). It is possible that higher parasitism in reproductive individuals will reduce their fitness, thereby producing lactation’s fitness cost – and, by extension, gestation’s lack of fitness cost – in this system (Clutton-Brock et al., 1989; Froy et al., 2016; Harshman & Zera, 2007; Williams, 1966). If exposure is determining parasitism and parasitism is reducing fitness, we would expect that parasite-mediated life history tradeoffs would be exacerbated in years and areas of high deer density, as more deer will translate to higher levels of pasture contamination (Wilson et al., 2004). Future investigations in this system could address the hypothesized role of parasite exposure and foraging behaviour in reproductive tradeoffs, using available census data (Clutton-Brock et al., 1982; Froy et al., 2018) to examine how annual, seasonal and spatial variation in habitat use and deer density correlate with environmental larval counts, parasite intensity, and the severity of reproductive tradeoffs.

Reproductive tradeoffs are a potential driver of seasonal dynamics of immunity and parasitism, in which periodic reproduction-associated relaxation of immunity leads to increased parasitism (Martin, Weil, & Nelson, 2008). Our results do not support this mechanism for several reasons: all status categories exhibited seasonality of antibodies, strongyles, and *E. cervi* rather than only reproductive individuals; reproductive increases in parasitism were not linked to lower immunity; and immunity did not correlate with resource availability, being highest in April, when the deer are in poor condition, having just survived the winter. In fact, antibody levels and strongyle counts correlated positively across seasons despite their negative correlation among individuals. This suggests that seasonality of propagule output is adaptive for helminths, facilitating highest transmission when environmental conditions are favourable and immunologically naïve calves are present, and leading to seasonal upregulation of immunity in warmer months to combat increased exposure (Møller, Erritzøe, & Saino, 2003; Wilson et al., 2004).

This study describes unexpected and complex interrelationships between different components of mammalian reproduction, immunity, and parasitism in the wild. We suggest that classical resource allocation mechanisms which are often hypothesised to underlie tradeoffs with immunity (e.g. Sheldon & Verhulst 1996; Deerenberg *et al*. 1997; French *et al*. 2007) are insufficient to explain many of the patterns seen in wild mammals, corroborating findings in other taxa (Stahlschmidt et al., 2013; Svensson et al., 1998). As such, studies examining such tradeoffs in mammals should consider mechanistic links between reproduction and immunity, resource acquisition limitations, and exposure components of parasitism, particularly by quantifying both immunity and parasitism simultaneously (Bradley & Jackson, 2008; Graham et al., 2011). The potential complexity of such interrelationships may contribute to the relative rarity of conclusive evidence for reproduction-immunity-parasitism tradeoffs in mammals.

## Acknowledgments

The long term red deer study is funded by the Natural Environment Research Council (grant number NE/L00688X/1), as is GFA’s PhD studentship through the E3 Doctoral Training Partnership (grant number NE/L002558/1). FK receives funding from the Scottish Government, RESAS, Strategic Research Programmes 2016-21. We thank Scottish Natural Heritage for permission to work on the Isle of Rum NNR and for the support of the reserve management team on the island. Thanks to Dave McBean and Gillian Mitchell at the Moredun Research Institute for their help with parasitological methods. The *Teladorsagia circumcincta* antigen was received from Moredun Research Institute, and was prepared by David Bartley, Alison Morrison, Leigh Andrews, David Frew and Tom McNeilly. Thanks also to Eryn Macfarlane and Adam Hayward for their helpful comments on the manuscript, and to Olly Gibb and all field assistants for their help in sample collection.

## Supplementary information

### Section One: Table SI1

**Table SI1:** model sample sizes and repeatabilities (95% credibility intervals in brackets).

| Factor | Prevalence (%) | Model Sample Sizes |  |  | Repeatability |
| --- | --- | --- | --- | --- | --- |
|  |  | Full | Pregnancy | Calving Traits |  |
| **Strongyles** | 76 | 835 | 518 | 571 | 0.21 (0.16-0.28) |
| ***F. hepatica*** | 33 | 824 | 517 | 571 | 0.11 (0.06-0.17) |
| ***E. cervi*** | 95 | 833 | 518 | 571 | 0.39 (0.34-0.45) |
| **Total IgA** |  | 796 | 499 | 550 | 0 (0-0.08) |
| **Anti-Tc IgA** |  | 796 | 497 | 547 | 0.25 (0.17-0.32) |

### Section Two: Model output

This section includes model outputs for the fixed effects of all models we ran. We first include the main models reported in the study (Figure SI1). We then compare these results with a set of modifications that we investigated.

The next (Figure SI2) displays the full models including a season by status interaction to display the way this affected the estimates, and to demonstrate the seasonal effects. Generally, including a season by status interaction did not improve the model fit or change our findings. The exception for this is the strongyle model (delta DIC = −3.79). There was, however, a general trend for the differences between status categories to decrease over the autumn and spring seasons as can be seen in Figure 3 in the main text.

Figure SI3 displays the effects from pregnancy models when we either included or removed pregnancy as a fixed effect, to investigate whether this affected the estimates of each status category’s effect. Inclusion of the pregnancy term slightly reduced the significance of the “Calf Survived” effect in the strongyle model, and increased the effects of “Calf Survived” in both the total IgA and anti-Tc IgA models. It also improved the fit of the total IgA model (delta DIC = −3.71). Otherwise, pregnancy had little effect.

Figure SI4 displays the results from the full models again, compared with the results from the multivariate models. The models were extremely similar, with only small changes in effect sizes and significance. This validates our use of the model to derive phenotypic correlations.

**Figure SI1:**
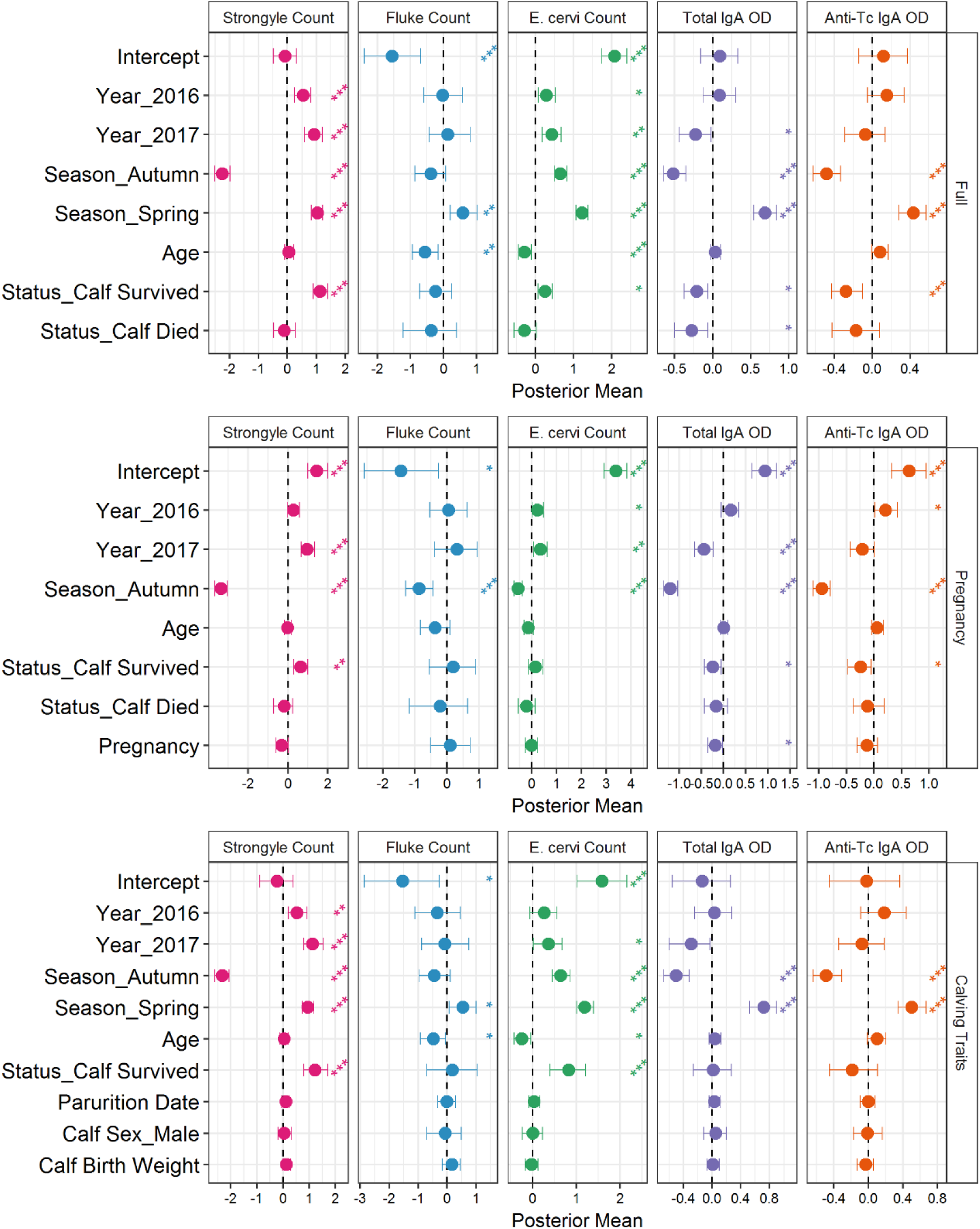
Effect size estimates from the three model sets (full dataset, pregnancy models and calf traits models). Effect sizes for categorical variables denote differences from the first (absent) category of each, contained in the intercept. Effect sizes for continuous variables represent the change associated with a change of one standard deviation of the variable in question. Points and error bars represent model estimates and 95% credibility estimates for each of the five full models. Asterisks indicate the significance of variables: ***, ** and * indicate P<0.001, P<0.01 and P<0.05 respectively.

**Figure SI2:**
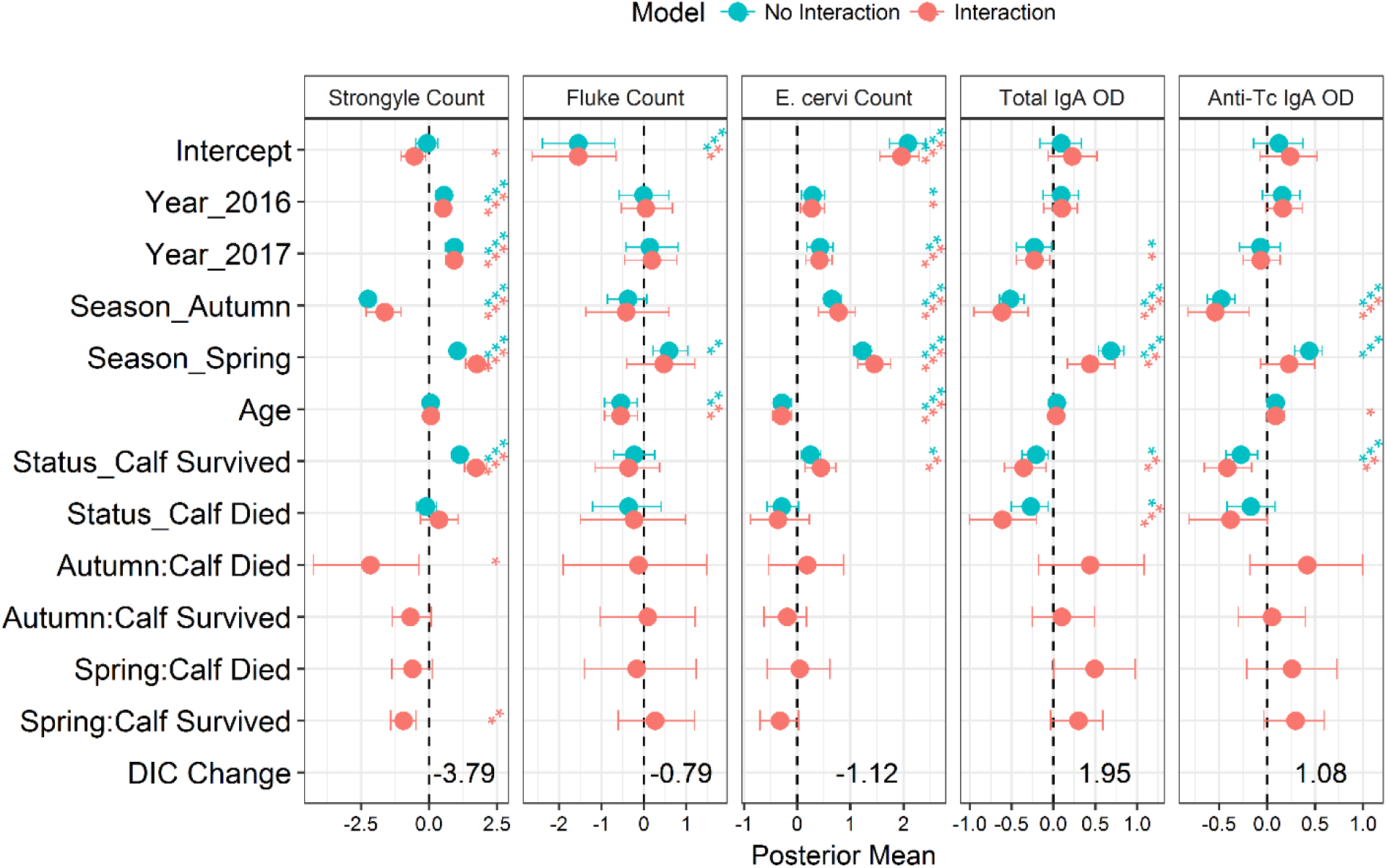
Comparison between outputs from full models excluding and including a season by status interaction. Points and error bars represent model estimates and 95% credibility estimates for each of the five full models, with and without interactions. Effect sizes for categorical variables denote differences from the first (absent) category of each, contained in the intercept. Effect sizes for continuous variables represent the change associated with a change of one standard deviation of the variable in question. Asterisks indicate the significance of variables: ***, ** and * indicate P<0.001, P<0.01 and P<0.05 respectively. DIC Change represents the change in DIC that occurred when an interaction was included. Including an interaction did not have a substantial effect on most of the original estimates, and only the fit of the strongyle model was significantly improved by its inclusion.

**Figure SI3:**
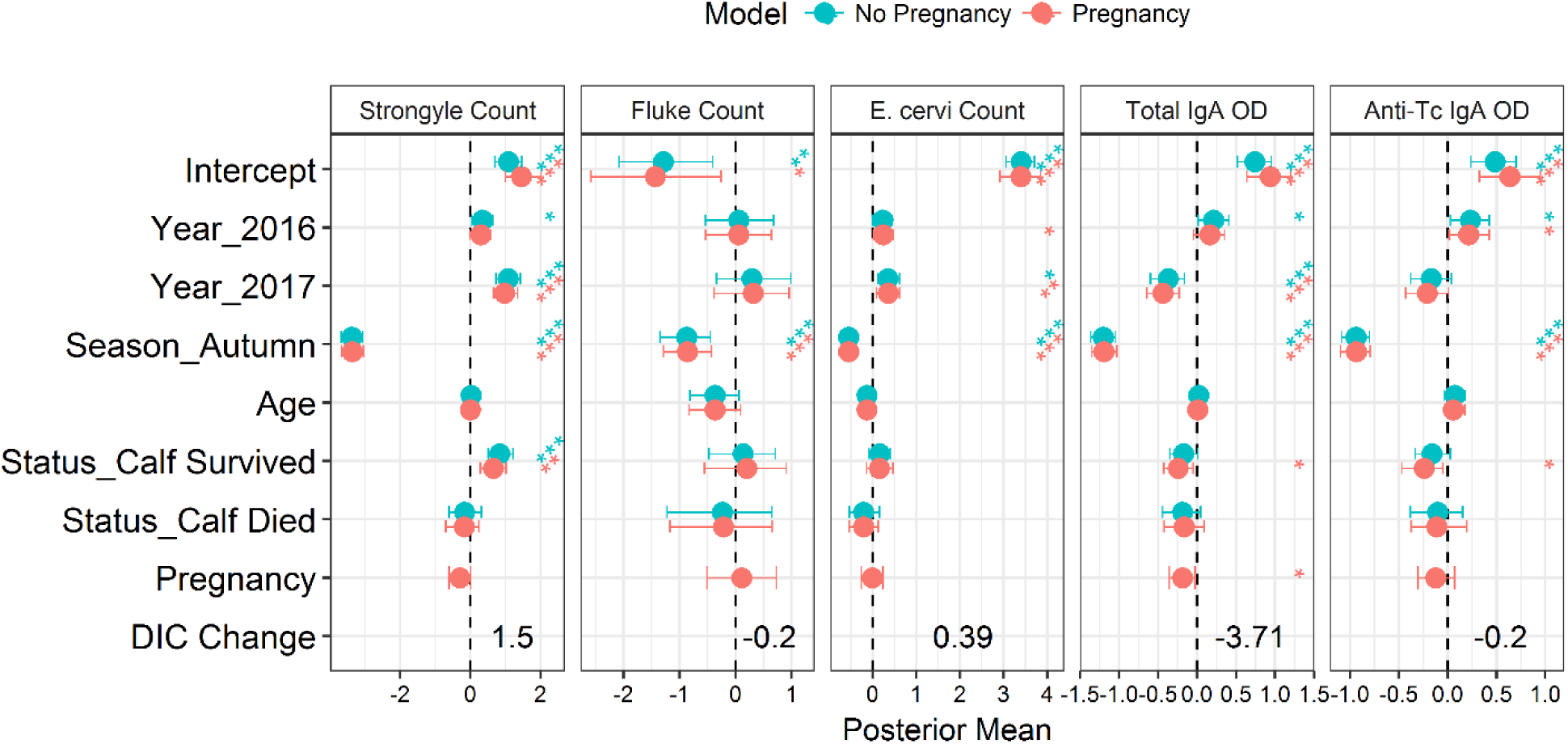
Comparison between outputs from pregnancy models, excluding and including pregnancy as a covariate to investigate whether this changed the model estimates for the reproductive status categories. Points and error bars represent model estimates and 95% credibility estimates for each of the five full models, without and with pregnancy as a covariate. Effect sizes for categorical variables denote differences from the first (absent) category of each, contained in the intercept. Effect sizes for continuous variables represent the change associated with a change of one standard deviation of the variable in question. Asterisks indicate the significance of variables: ***, ** and * indicate P<0.001, P<0.01 and P<0.05 respectively. DIC Change represents the change in DIC that occurred when pregnancy was included. Including pregnancy as a covariate slightly decreased the effect size of “Calf Survived” for strongyles and increased it for total IgA and anti-Tc IgA, but otherwise had little effect on the estimates. Pregnancy also significantly reduced total IgA levels and improved the total IgA model fit.

**Figure SI4:**
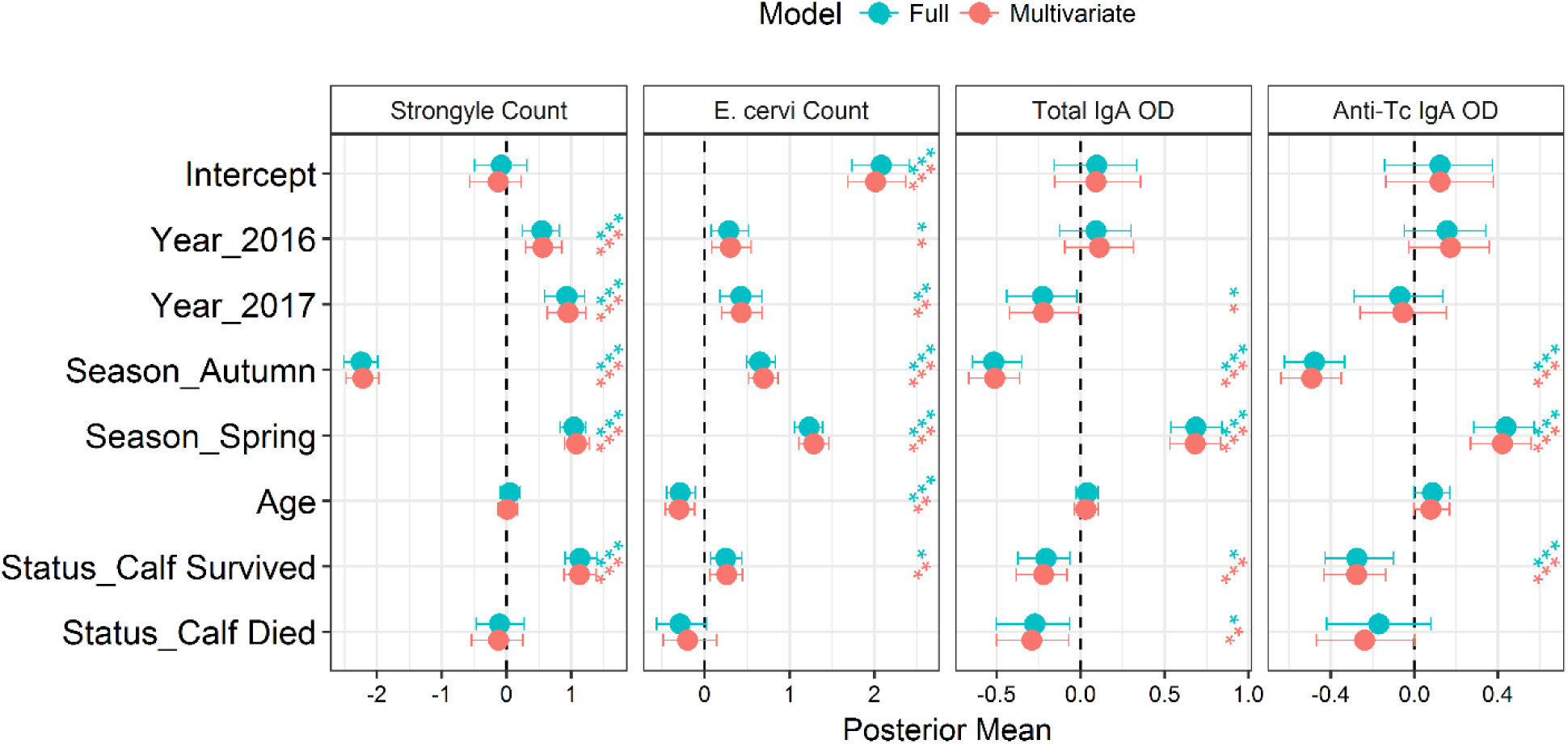
Comparison of the fixed effect estimates from the full models and the multivariate model. Points and error bars represent model estimates and 95% credibility estimates for each of the five full models and the equivalent fixed effects in the multivariate model. Effect sizes for categorical variables denote differences from the first (absent) category of each, contained in the intercept. Effect sizes for continuous variables represent the change associated with a change of one standard deviation of the variable in question. Asterisks indicate the significance of variables: ***, ** and * indicate P<0.001, P<0.01 and P<0.05 respectively.

### Section Three: Additional Figures

This section contains some figures displaying patterns in the data which are of interest. This includes the effects of faecal collection variables on antibody levels, age effects and correlations between response variables.

**Figure SI5:**
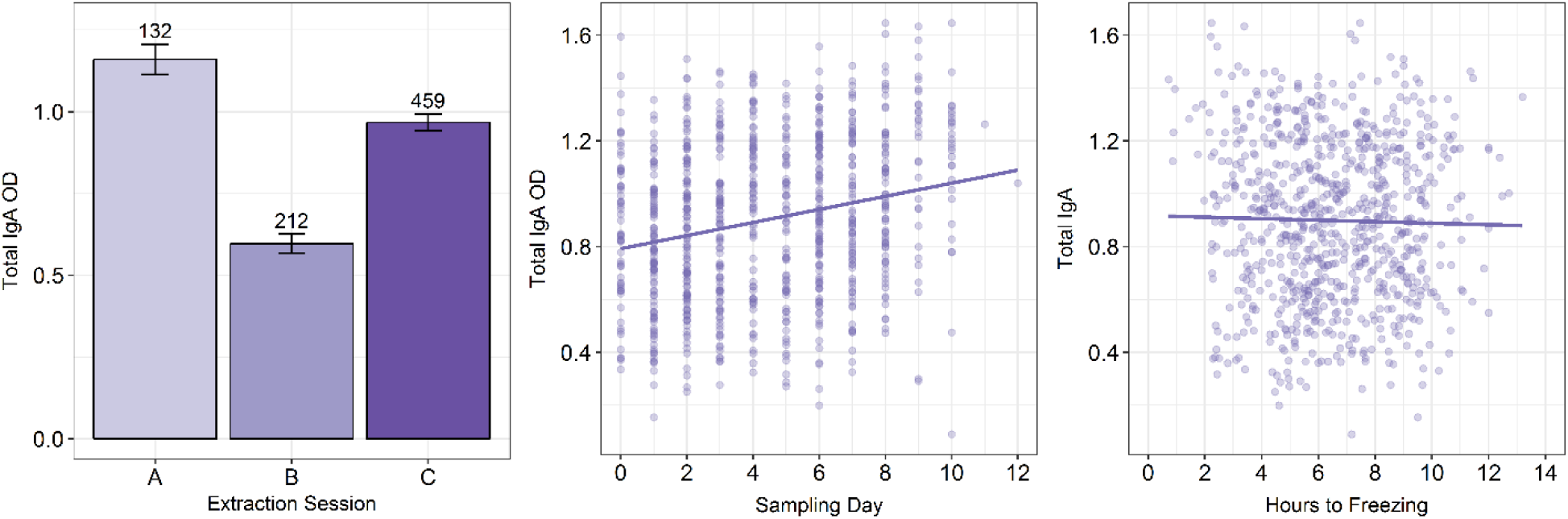
Impact of faecal collection factors on total IgA level (extraction session, day of collection and hours to freezing). Y axes for figures B and C have been square root transformed.

**Figure SI6:**
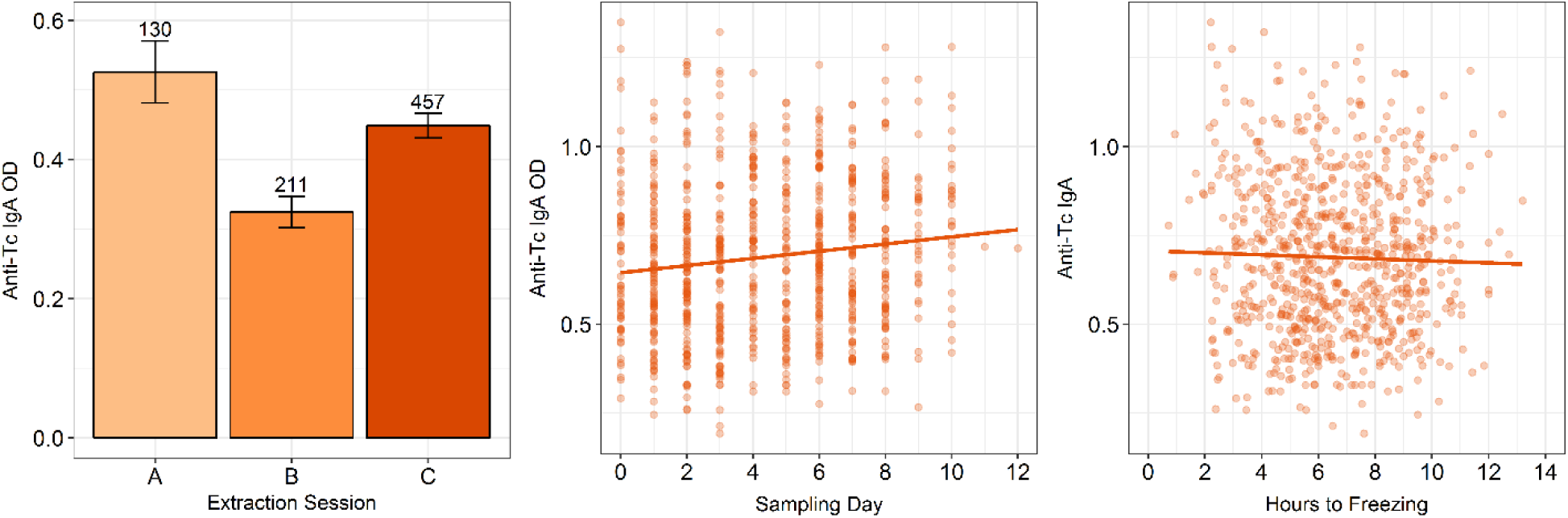
Impact of faecal collection factors on anti-Tc IgA level (extraction session, day of collection and hours to freezing). Y axes for figures B and C have been cube root transformed.

**Figure SI7:**
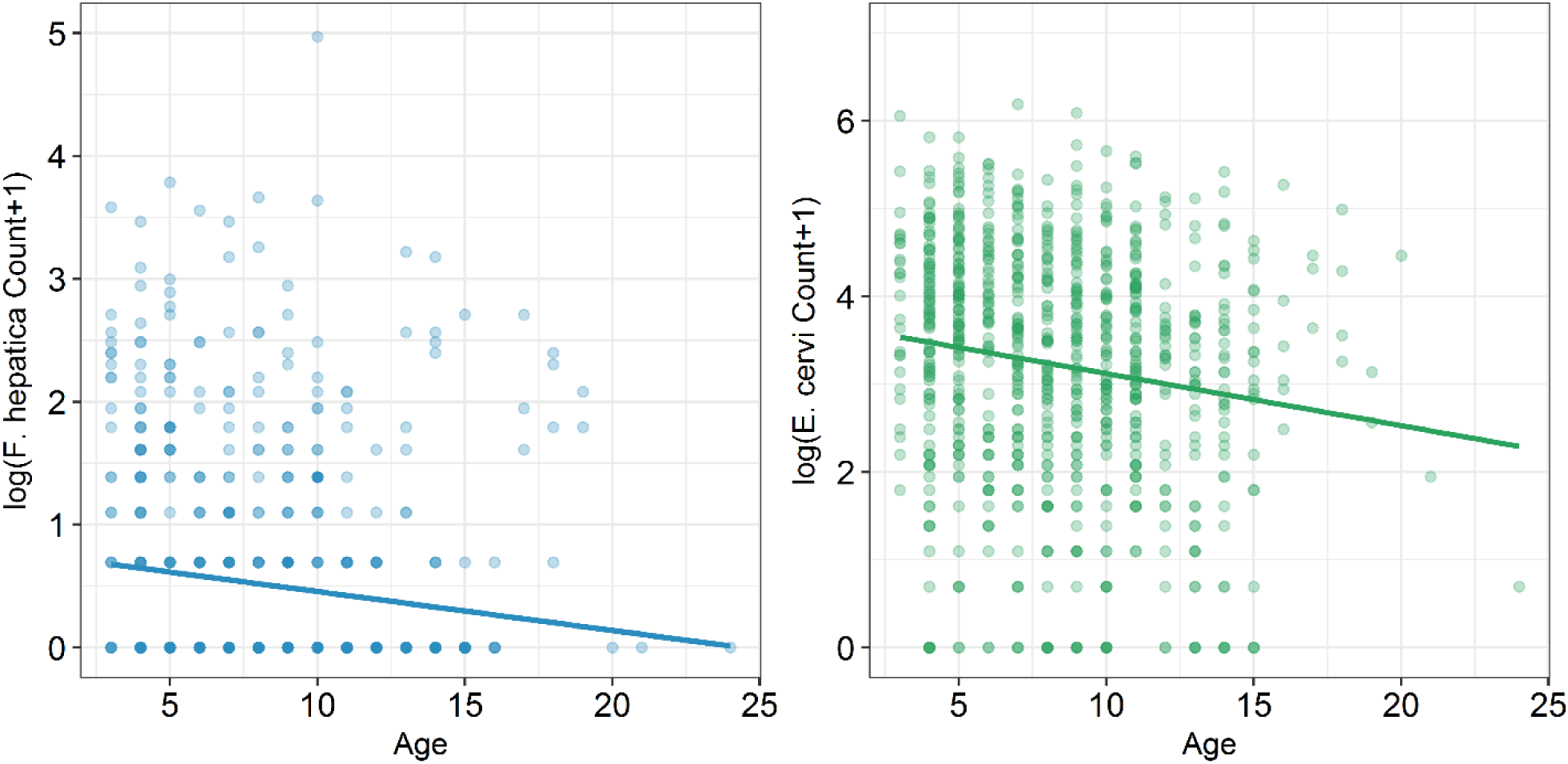
Age trends in parasite counts using the raw correlations (*F. hepatica* and *E. cervi*). Y axes have been log(count+1)-transformed.

**Figure SI8:**
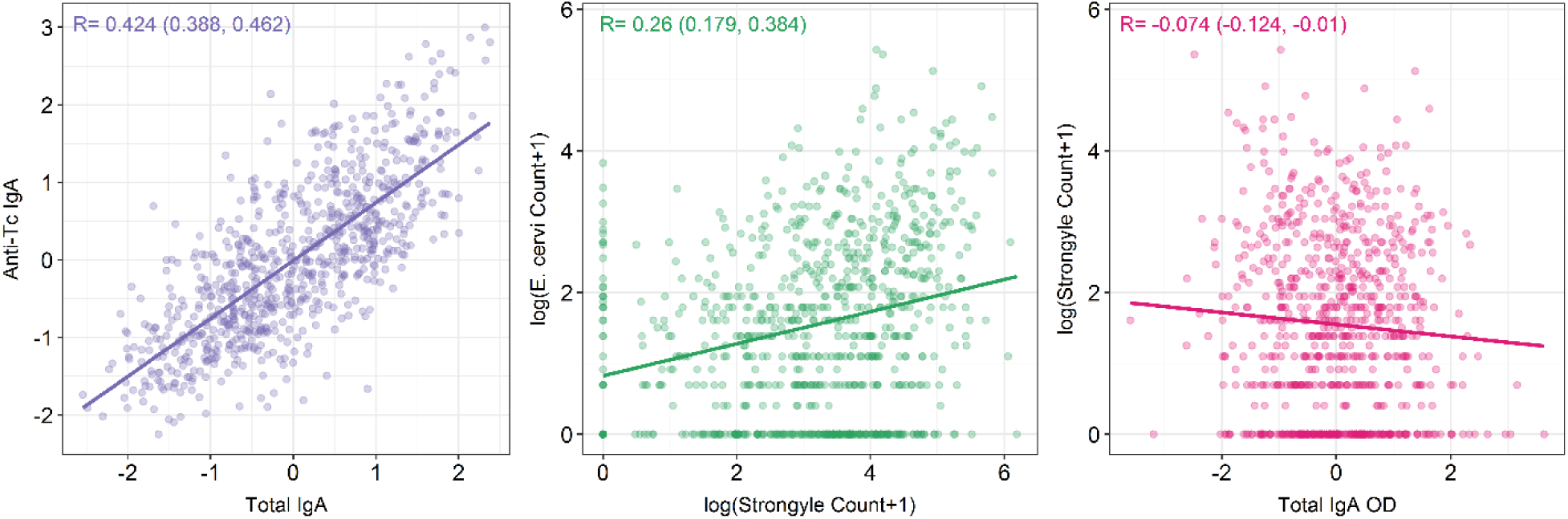
Correlations between response variables (Total IgA and anti-Tc IgA; Strongyles and E. cervi; Total IgA and Strongyles). Model-derived phenotypic correlations (R_p_) are included, with 95% credibility intervals. Both antibody measures are based on the residuals from a model including extraction session, day of collection and hours to freezing, with transformed antibody OD as response variable(square root for the total IgA and cube root for anti-Tc IgA). In the strongyle figure total IgA was scaled within each sampling trip to have a mean of 0 and a standard deviation of 1 to avoid a positive correlation arising from shared seasonal effects.

**Author contributions**
GFA collected the samples, conducted labwork, analysed the data, and drafted the manuscript; KW designed and helped to carry out the ELISAs; RK carried out some antibody extractions and ELISAs; SM and AM helped with sample collection; FK, DN, JP offered comments on methodology and theory throughout and helped draft the manuscript.

